# Cryo-EM structure of the transcription termination factor Rho from *Mycobacterium tuberculosis* reveals mechanism of resistance to bicyclomycin

**DOI:** 10.1101/2021.06.25.449899

**Authors:** Emmanuel Saridakis, Rishi Vishwakarma, Josephine Lai-Kee-Him, Kevin Martin, Isabelle Simon, Martin Cohen-Gonsaud, Franck Coste, Patrick Bron, Emmanuel Margeat, Marc Boudvillain

**Affiliations:** Institute of Nanoscience and Nanotechnology, NCSR “Demokritos”, Ag. Paraskevi 15310, Athens, Greece; Le Studium Loire Valley Institute for Advanced Studies, Orléans, France; Centre de Biologie Structurale, INSERM, CNRS, Université de Montpellier, 34090 Montpellier, France; Department of Biochemistry and Molecular Biology, The Pennsylvania State University, University Park, State College PA-16802, USA; ED 549, Santé, Sciences Biologiques & Chimie du Vivant, Université d’Orléans, France; Centre de Biophysique Moléculaire, CNRS UPR4301, rue Charles Sadron, 45071 Orléans cedex 2, France; affiliated with Université d’Orléans

## Abstract

The bacterial Rho factor is a ring-shaped motor triggering genome-wide transcription termination and R-loop dissociation. Rho is essential in many species, including in *Mycobacterium tuberculosis* where *rho* gene inactivation leads to rapid death. Yet, the *M. tuberculosis* Rho [_Mtb_Rho] factor displays poor NTPase and helicase activities, and resistance to the natural Rho inhibitor bicyclomycin [BCM] that remain unexplained. Here, we address these unusual features by solving the cryo-EM structure of _Mtb_Rho at 3.3 Å resolution, providing a new framework for future antibiotic development. The _Mtb_Rho hexamer is poised into a pre-catalytic, open-ringed state wherein specific contacts stabilize ATP in intersubunit ATPase pockets, thereby explaining the cofactor preference of _Mtb_Rho. We reveal a leucine-to-methionine substitution that creates a steric bulk in BCM binding cavities near the positions of ATP γ-phosphates, and confers resistance to BCM at the expense of motor efficiency.

## INTRODUCTION

Tuberculosis is a major global health concern, each year killing ~1.5 million people worldwide. Multi-resistant strains of *Mycobacterium tuberculosis* −the causative agent of tuberculosis-arise at an alarming rate and there is an urgent need to better understand the mechanisms of drug resistance and to develop alternative therapeutic strategies against *M. tuberculosis*^1^.

Transcription termination factor Rho is a central component of gene regulation in bacteria. It is essential in many Gram-negative species and in high G+C Gram-positive Actinobacteria such as *M. tuberculosis*^2^ or *Micrococcus luteus*^3^. Since Rho has no structural homologs in eukaryotes, it is an attractive target candidate for the development of new antibiotics^4^.

Some Actinobacteria of the *Streptomyces* genus produce a natural Rho inhibitor called Bicyclomycin [BCM]. BCM is effective against various Gram-negative pathogens but is inactive against most Gram-positive species, including *M. tuberculosis*^5^. One notable exception is *M. luteus*, whose growth is inhibited by BCM^3^. Accordingly, BCM strongly inhibits the *in vitro* enzymatic activity of *M. luteus* Rho (_Mic_Rho)^3^ but hardly affects *M. tuberculosis* Rho (_Mtb_Rho)^6^. There is currently no rational explanation for this difference.

Our understanding of the mechanisms of Rho-dependent transcription termination (RDTT) and inhibition by BCM stems mostly from studies of *Escherichia coli* Rho (_Ec_Rho). The _Ec_Rho prototype is a ring-shaped, hexameric protein motor that dissociates transcription elongation complexes (TECs) in an RNA- and ATP-dependent manner^7–9^. Activation of _Ec_Rho is triggered by binding to a C>G sequence-skewed and poorly structured *Rut* (Rho utilization) site in the nascent transcript and by allosteric closure of the _Ec_Rho ring around the RNA chain^7–10^. Once activated, the _Ec_Rho ring can hydrolyze ATP, translocate RNA, unwind RNA:DNA duplexes, and disrupt TECs^11–14^. Recent work supports a model where the EcRho motor first binds RNAP and allosterically destabilizes the TEC from this sitting position once it senses a *Rut* site in the emerging transcript^15–17^.

The six N-terminal domains (NTDs) of _Ec_Rho form a crown-like Primary binding site (PBS) on one face of the hexamer ring^18^. This composite PBS includes a YC-binding pocket (Y being a C or U residue) on each _Ec_Rho protomer that contributes to the specific recognition of transcript *Rut* sites^18,19^. The NTD also carries the residues contacting RNA polymerase (RNAP) identified in recent cryoEM structures of the _Ec_Rho:RNAP complex^15,16^.

Despite these important roles, the NTD is not highly conserved and often contains large N-terminal insertion domains (NIDs), in particular in high G+C Actinobacteria (**Figure S1**)^4^. In _Mic_Rho and _Mtb_Rho, these inserts increase the affinity for RNA, allow productive interaction with structured transcripts, and promote RDTT at promoter-proximal sites that the NID-less _Ec_Rho is unable to use^6,20^. Indels also interrupt the PBS sequences of MicRho and MtbRho (**Figure S1**), which may be further indications of a *Rut*/RNA sensing mechanism deviating from the EcRho paradigm.

The much more conserved C-terminal domain (CTD) of _Ec_Rho is responsible for intersubunit cohesion and ATP-dependent RNA translocation^10,21^. The CTD notably carries the Walker A/B motifs forming ATPase pockets at subunit interfaces and the “catalytic Glu’, ‘Arg valve’, and ‘Arg finger’ residues required for catalysis of ATP hydrolysis. These residues are highly conserved in phylo-divergent Rho factors, including in _Mtb_Rho (**Figure S1**)^4^.

The CTD also contains the secondary binding site (SBS) Q- and R-loop motifs that translocate RNA through the EcRho ring as a function of the chemical state of the ATPase pockets^10,21^. A K→T mutation in the R-loop (Lys326 in _Ec_Rho to Thr501 in _Mtb_Rho) slightly weakens enzymatic activity^6^. The CTD also carries the side chains that form an interaction pocket for BCM near each ATP binding site^22^. Binding of BCM to these pockets locks _Ec_Rho in the open ring conformation, thereby preventing enzymatic activation^23^. The BCM-binding side-chains are strictly conserved in _Mtb_Rho (**Figure S1**) and thus cannot account for its resistance to BCM.

To elucidate the origin of this resistance to BCM and to better comprehend the evolutionary specifics of RDTT in *M. tuberculosis*, we solved the structure of _Mtb_Rho using cryo-EM single-particle analysis. Initial attempts by X-ray crystallography proved fruitless (see supplementary information) while cryo-EM led to a 3.3 Å resolution map. We show that _Mtb_Rho can adopt an open, ring-shaped hexamer conformation that mimics that observed for _Ec_Rho^18^. The NIDs are not resolved in the _Mtb_Rho hexamer structure, consistent with predicted intrinsically disordered features. We identify a leucine-to-methionine substitution in _Mtb_Rho that creates a steric bulk in the cavity where BCM normally binds. We show that this Leu→Met mutation is a taxa-specific evolutionary feature that alone is sufficient to account for _Mtb_Rho resistance to BCM.

## RESULTS AND DISCUSSION

### High-resolution reconstruction of the _Mtb_Rho complex

Freshly purified _Mtb_Rho mostly forms monodisperse hexamers in presence of Mg-ATP and dC_20_ ligands, as determined from SEC-MALS experiments (**Figure S2A&B**). Images of negatively stained _Mtb_Rho complexes reveal the hexameric organization of particles forming closed rings (**Figure S2C**) while these rings appear mostly open in conventional cryo-EM images (**Figure S2D**). The most suitable conditions for high-resolution image acquisition in terms of _Mtb_Rho particles distribution and orientation in ice in presence of Mg-ATP and dC_20_ ligands were obtained using Lacey grids. Upon processing 10,888 movies as described in **Figures S3** and **S4**, we were able to compute a cryo-EM map at 3.3 Å resolution and to confirm an open-ring organization of the complex resulting from the assembly of six _Mtb_Rho subunits (**Figure 1**). Local resolution mapped on the structure with the RELION package ranges from 3.1 to 4.8 Å (**Figure 1A-C**). A gradient of high-to-low resolution is apparent in the 3D map from the innermost _Mtb_Rho ring subunits (labelled with stars in **Figure 1B**) to subunits at the ring gap (labeled with black dots). This suggests some degree of flexibility among subunits, which was confirmed by a multibody analysis where the first main eigenvectors correspond to intersubunit twists (**Figure S5**). An atomic model of the _Mtb_Rho complex was built and refined based on the cryo-EM density map (**Table S1** and **Figure S6**) and is detailed below.

**Figure 1:**
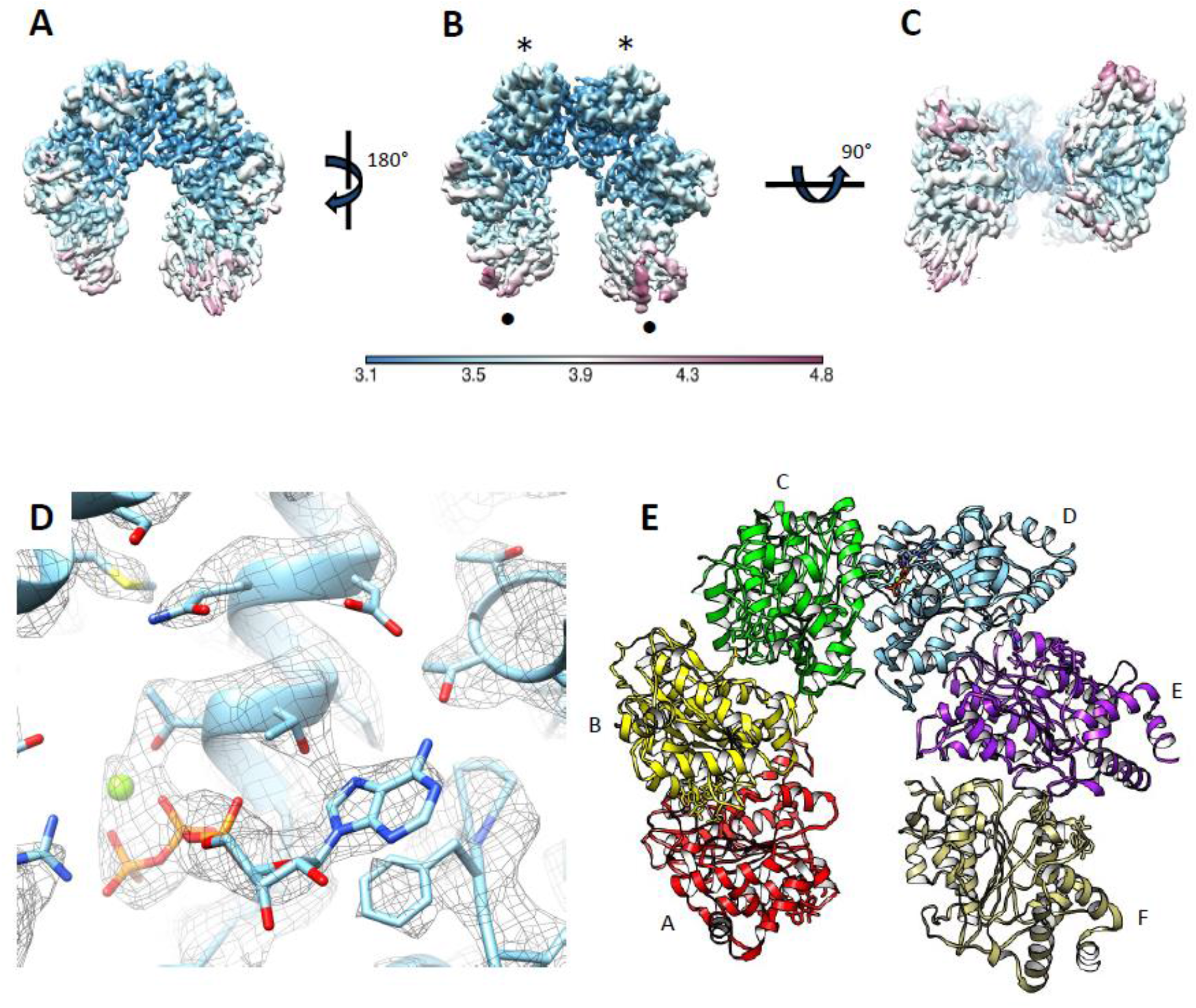
**(A, B, C)** The 3.3 Å cryoEM map of the _Mtb_Rho factor coloured by local resolution reveals an hexamer ring in an open corkscrew configuration. Local resolution estimates (color bar in Å) were calculated in RELION after the final refinement. The first ~220 N-terminal residues of each protomer are not resolved. **(D)** Representative density features fitted into the 3.3 Å cryoEM-based atomic model of the _Mtb_Rho hexamer (subunit D). Cryo-EM densities (grey mesh) are superimposed on the catalytic Mg^2+^ ion (apple green sphere), ATP cofactor, and surrounding _Mtb_Rho residues. **(E)** Refined PDB structure, obtained from the cryo-EM map. Subunits are labeled from A to F in clockwise order (looking at the CTD face of the _Mtb_Rho hexamer).

### The _Mtb_Rho hexamer adopts an open ring conformation

The _Mtb_Rho hexamer adopts an open, corkscrewed configuration (**Figure 1**) similar to the ones observed for _Ec_Rho (**Figure S7**) alone^18^ or in complex with _Ec_RNAP^15,16^. An open, corkscrewed hexamer was also proposed for RNA-free Rho from *Thermotoga maritima*^24^, supporting that this is a conserved and mechanistically important trait of the termination factor.

The N-termini, including NIDs, are not resolved in the structure of the _Mtb_Rho hexamer (**Figure 1**). This is consistent with the intrinsically disordered state of the NID as predicted by the XtalPred server^25^. The dC_20_ ligand is also not resolved despite its ability to bind and stabilize _Mtb_Rho, as determined by thermal shift assay^26^. It might be that dC_20_ interacts with NTD (or NID) but is not long enough to induce sufficient folding for the domain to be resolved. Alternatively, the NTD/NID may properly fold upon dC_20_ binding but adopt variable orientations with respect to the CTD due to a flexible NTD-CTD junction (**Figure 2A**). In this case, the dC_20_ ligand might be too short to bridge several NTDs/NIDs and restrict their movements. It is also possible that the NIDs substitute for the lineage-specific motifs that stabilize the _Ec_(Rho:RNAP) complex^15,16^ and become structurally organized only upon _Mtb_Rho interaction with RNAP^27^.

**Figure 2:**
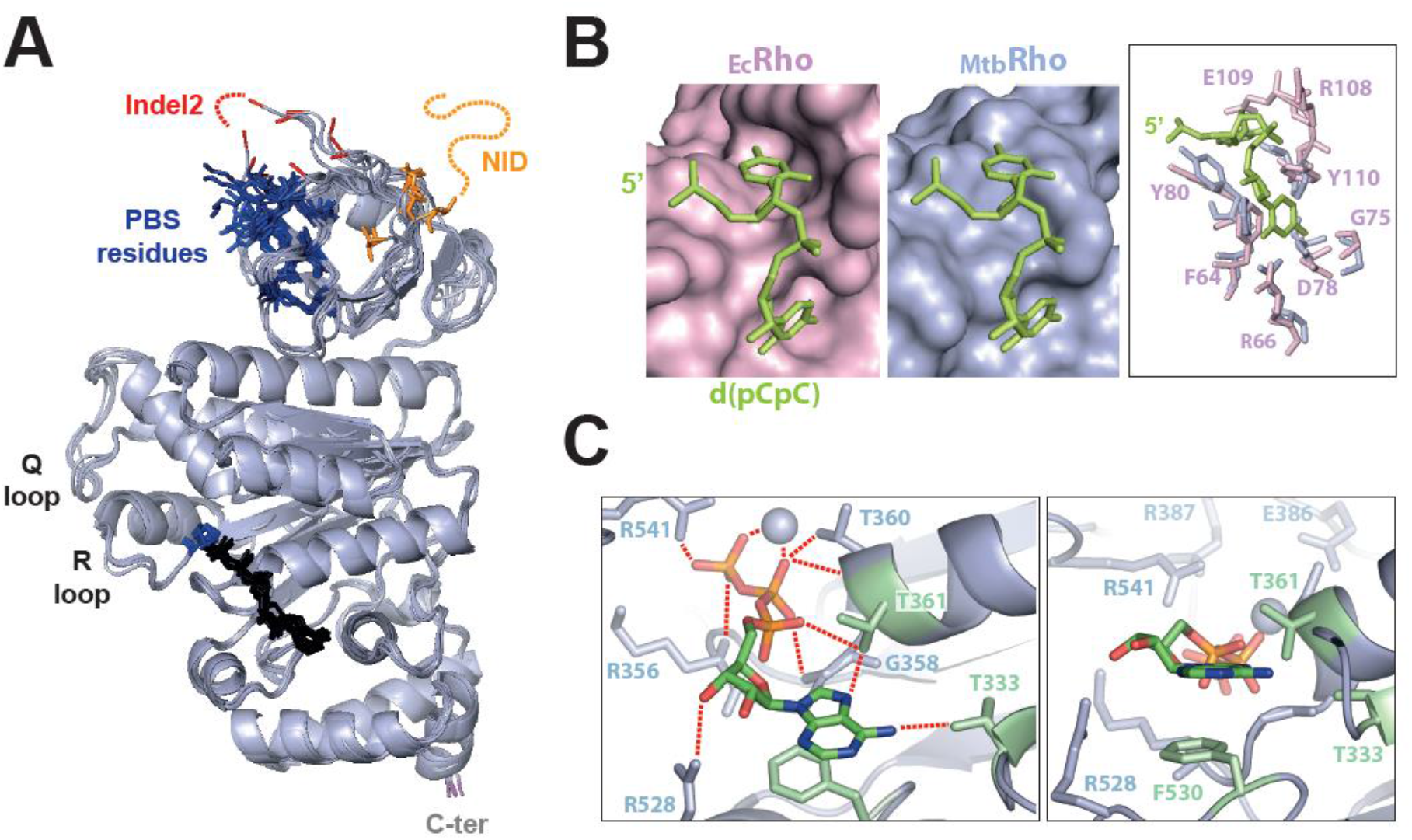
Organization of the ligands binding network in _MtbRho_. **(A)** Superposition of the six _Mtb_Rho protomers with the main RNA and ATP binding motifs highlighted. ATP ligands are shown in black with closeby Met495 side-chains in blue. Residues equivalent to the PBS residues of _Ec_Rho are also shown in blue. **(B)** Comparison of the PBS pockets of _Ec_Rho and _Mtb_Rho. A dC_2_ ligand (in green) has been modeled on _Mtb_Rho upon structural alignment with _Ec_Rho (PDB 1PV4). **(C)** Closeup views of the ATPase pocket at the best resolved interface (C/D) of _Mtb_Rho showing the network of interaction with the ATP ligand and catalytic Mg^2+^ ion (sphere).

PBS Glu280 and indel2 residues (**Figure S1**) are also not resolved while the other PBS side-chains adopt somewhat variable orientations in the _Mtb_Rho protomers (**Figure 2A**), possibly because of the lack of stabilizing nucleic acid ligand. As a result, the PBS pockets do not seem as deep or as defined as in _Ec_Rho^18^, particularly at the level of the 5’C-binding subsite (**Figure 2B**). Unfortunately, these structural features do not provide clues as to why _Mtb_Rho, but not _Ec_Rho, can bind and utilize structured RNA substrates efficiently^6,27,28^. Further work, e.g. with longer nucleic acid ligands, will be needed to address this specific question.

### Organization of the ATP-dependent allosteric network in _Mtb_Rho

The presence of a clear electron density in all the ATPase pockets of _Mtb_Rho reveals that ATP ligands are held by a network of interactions involving the phosphate, sugar, and adenine base moieties (**Figures 1D, 2A&C**). The interaction network also includes a catalytic Mg^2+^ ion coordinating β and γ-phosphate oxygens (**Figures 1D & 2C**). The adenine base is stacked on Phe530 while Thr333 and Thr361 form hydrogen bonds with the NH_2_ and N_7_ groups of adenine, respectively (**Figure 2C**). These H-bonds explain the marked preference of _Mtb_Rho for purine versus pyrimidine triphosphates^6^. Purine-specific interactions are not found in _Ec_Rho, which hydrolyzes the four NTPs with comparable efficiencies^29^. In this case, the NTP base is held in sandwich by an aromatic contact with _Ec_Phe355 (the equivalent of _Mtb_Phe530) and a methionine-aromatic contact with _Ec_Met186 (replaced by Thr361 in _Mtb_Rho) while _Ec_Thr158 (the equivalent of _Mtb_Thr333) is too far away (> 3.5 Å in all _Ec_Rho structures) to contribute significantly to the interaction (**Figure S8**)^10,18,21^.

Consistent with the open configuration of the _Mtb_Rho ring (**Figure 1**) and absence of activating RNA ligand, the ATPase pockets are in an unproductive state. A direct interaction with the arginine finger (R541) from the adjacent protomer somewhat shields the γ-phosphate of ATP (**Figures 2C** and **S8**). Furthermore, the arginine valve (R387) and catalytic glutamate (E386) residues are too distant from, respectively, the γ-phosphate and catalytic Mg^2+^ ion to catalyze ATP hydrolysis (**Figures 2C** and **S8**). By comparison, structures of the closed _Ec_Rho ring (i.e. the catalytically-competent state) display tighter subunit interfaces (**Figure S8**) where the catalytic Glu (E211) and Arg valve (R212) residues are adequately positioned to bind and polarize the catalytic Mg^2+^ ion and catalytic water molecule, respectively^10,21^. Tight contacts between the _Ec_Rho subunits also allow establishment of the allosteric communication network connecting the ATPase pocket to the RNA-binding SBS in the hexamer central channel (**Figure S9A**)^10^. Although all the network residues are conserved in _Mtb_Rho, the corkscrewed disposition of the subunits (**Figure S7**) prevents formation of the network contacts (**Figure S9A**). The allosteric network is also not formed in the ‘pre-catalytic’ open ring structure of _Ec_Rho^18^.

The R- and Q-loops of _Mtb_Rho are connected by a H-bond between the hydroxyl group of Thr501 and the carbonyl of Gly462 (**Figure S9B**). Such an interaction is not observed in _Ec_Rho where the Lys326 side-chain (the equivalent of _Mtb_Thr501) instead extends into the central channel (**Figure S9B**) as part of the RNA-binding SBS^10,21^. The H-bonded _Mtb_Thr501 side-chain cannot play this role in _Mtb_Rho (**Figure S9B**), suggesting that the SBS is restricted to Q-loop residues and explaining why a Thr→Lys mutation at position 501 stimulates the enzymatic activity of _Mtb_Rho^6^.

### A single Leu→Met mutation creates a steric constraint in the BCM binding pocket of _Mtb_Rho

The totality of residues identified as directly interacting with BCM in the crystal structure of the _Ec_Rho-BCM complex^22^ are strictly conserved in _Mtb_Rho (**Figure S1**)^4^. However, close inspection of the BCM-binding cavity in our _Mtb_Rho structure revealed that it contains a methionine (Met495) instead of a leucine (Leu320) in _Ec_Rho. This Leu→Met substitution creates a bulk in an otherwise structurally comparable cavity (**Figure 3A**), which could penalize BCM binding by steric clash. The hindrance could be with BCM itself or with the _Mtb_Lys359 side-chain from the Walker A motif (aka P-loop). In _Ec_Rho, the corresponding lysine (Lys184) undertakes a significant conformational change upon BCM binding. This movement may be impaired by the bulky Met495 neighbor in _Mtb_Rho (**Figure S10A**).

**Figure 3:**
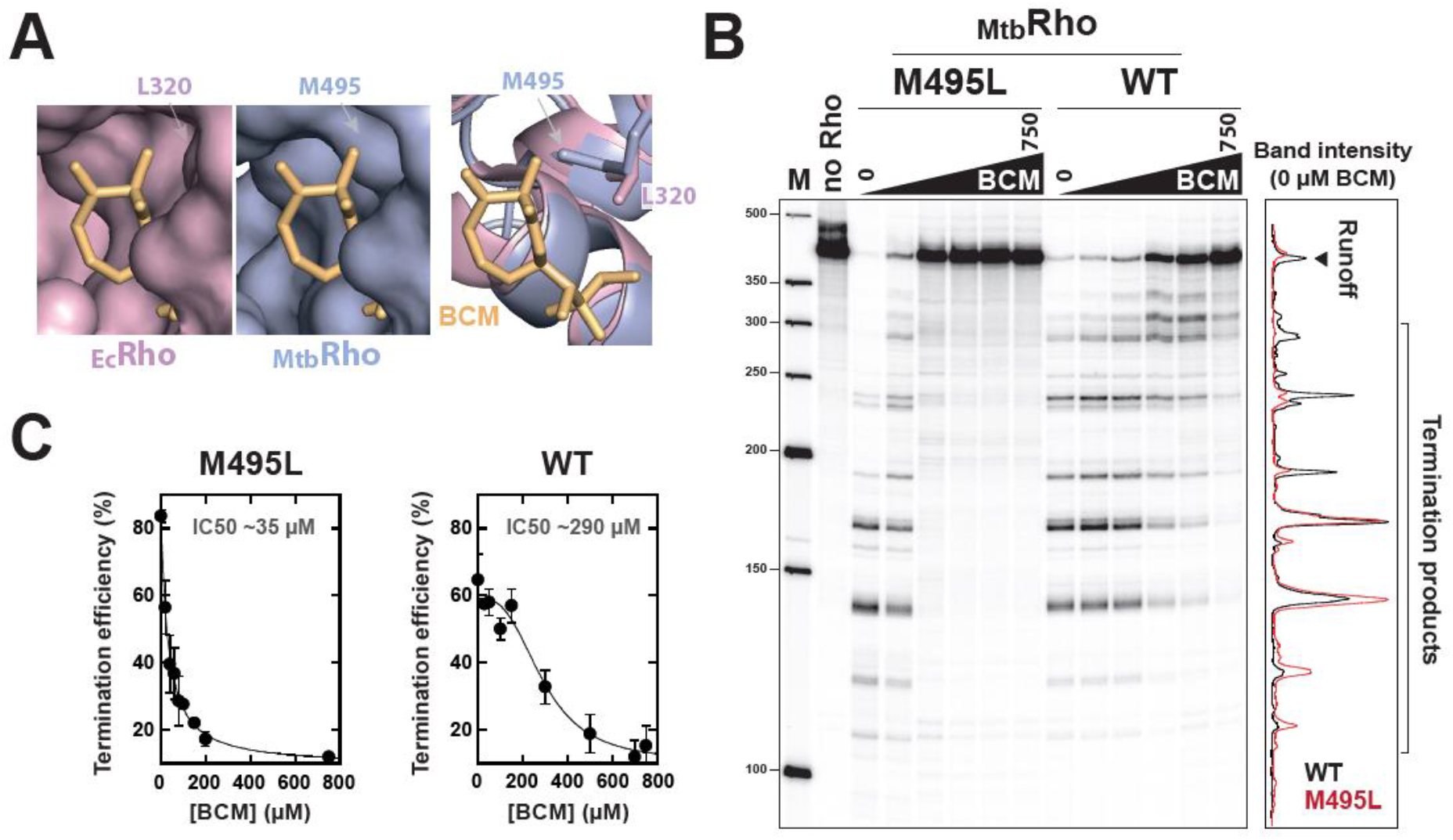
The _Mtb_Met495 side-chain confers BCM resistance to _Mtb_Rho. **(A)** The BCM binding pockets of _Ec_Rho (PDB 1PV4) and _Mtb_Rho (this work). BCM has been fitted into the pockets by structural alignment with BCM-bound _Ec_Rho (PDB 1XPO). **(B)** The M495L mutation sensitizes _Mtb_Rho to BCM. A representative PAGE gel shows the effects of increasing concentrations of BCM (0-750 μM) on the transcription termination activities of the WT _Mtb_Rho and M495L mutant. The graph compares the gel lane profiles for the WT (black) and M495L (red) proteins in absence of BCM (lanes 0). **(C)** IC50 graphs deduced from two independent transcription termination experiments.

To test this ‘steric clash’ hypothesis, we compared the transcription termination activities of WT _Mtb_Rho and its M495L mutant derivative using *E. coli* RNAP and a DNA template encoding the Rho-dependent λtR1 terminator. With this *in vitro* heterologous system, WT _Mtb_Rho triggers efficient transcription termination, starting at promoter-proximal sites along the DNA template (**Figure 3B**), as described previously^6^. High concentrations of BCM are required to perturb the termination activity of WT _Mtb_Rho (**Figure 3B**)^6^. The M495L mutant is also a very efficient termination factor but its activity is readily inhibited by ~10 times lower concentrations of BCM (**Figure 3B&C**). Under the same experimental conditions, the IC50 values previously measured for the BCM-sensitive WT _Ec_Rho^27^ and here for the _Mtb_Rho M495L mutant (**Figure 3C**) are similar (~40 and ~35 μM, respectively). These data thus identify the Met495 side-chain as the main structural determinant of _Mtb_Rho resistance to BCM.

A Leu→Met mutation is found at the same position in the Rho sequences of various Actinobacteria of the Corynobacteriales order (**Figure S11**). This is notably the case for *Mycobacterium cholonae*, a bacterium that contains the cluster of genes responsible for BCM biosynthesis^30^. The Leu→Met mutation may thus render *M. cholonae* immune to its own BCM production. However, this protective mechanism does not appear to be widespread. Indeed, _Ec_Leu320 is strictly conserved in the Rho sequences of other bacteria bearing the BCM biosynthesis gene cluster (**Figure S11**), including the *Streptomyces* species known to produce BCM^31^.

Apart from Actinobacteria, we retrieved the Leu→Met mutation in two closely related insect endosymbionts from the Gram-negative Bacteroidetes phylum (**Figure S11**). This finding is surprising given the endosymbiont lifestyle, which itself should provide protection against BCM exposure. Overall, the Leu→Met mutation is rare and the only side-chain change observed at this position within the ~1300 representative Rho sequences compiled previously from multiple taxa^4^.

### An evolutionary tradeoff between enzymatic efficiency and BCM resistance

An unexpected effect of the M495L mutation is the stimulation of transcription termination when compared to WT _Mtb_Rho (~80% versus ~60% apparent efficiency in absence of BCM; **Figure 3C**). Early termination is also favored with the M495L mutant (**Figure 3B,** graph), suggesting that the Met495 side-chain reduces the efficiency of the _Mtb_Rho motor.

To confirm this idea, we compared the helicase activities of M495L and WT _Mtb_Rho. As described previously, WT _Mtb_Rho is unable to unwind long RNA:DNA hybrids (**Figure 4A**)^6^. By contrast, the M495L mutant readily unwinds a model RNA:DNA hybrid (57 base pairs), albeit at a slower rate than the _Ec_Rho control (**Figure 4A**). This residual deficiency may be due to the R-loop Thr501 side-chain (versus Lys326 in _Ec_Rho), which also reduces the enzymatic efficiency of _Mtb_Rho^6^. Alternatively, The *Rut* site within the model RNA:DNA substrate may be more adapted to _Ec_Rho than to _Mtb_Rho requirements.

**Figure 4:**
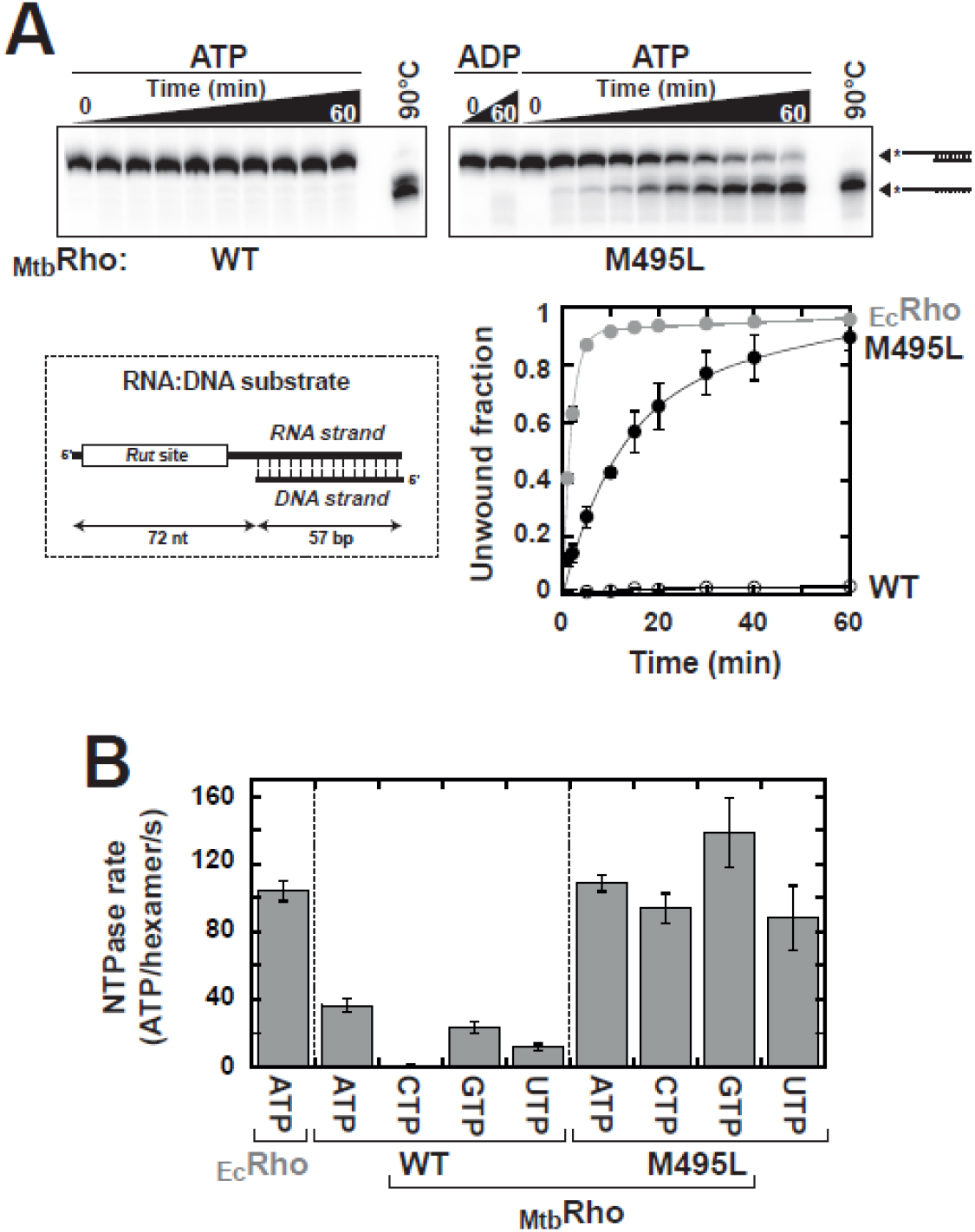
The Met495 side-chain impairs the activity of the _Mtb_Rho motor. The M495L mutation stimulates the RNA-DNA duplex unwinding **(A)** and NTP hydrolysis **(B)** activities of _Mtb_Rho. A loss in NTP selectivity is observed for the M495L mutant as compared to WT _Mtb_Rho.

Notwithstanding, in the presence of an excess of the model poly[rC] substrate, the M495L mutant hydrolyses ATP at a steady-state rate similar to that measured for _Ec_Rho and about 3 times faster than WT _Mtb_Rho (**Figure 4B**). Intriguingly, M495L hydrolyses the four rNTPs with comparable efficiencies (**Figure 4B**) whereas WT _Mtb_Rho has a marked preference for purine vs pyrimidine triphosphates^6^.

As mentioned above, pyrimidine triphosphates are probably less tightly bound in the ATPase pockets of _Mtb_Rho than in the pockets of _Ec_Rho due to a Met→Thr substitution (_Mtb_Thr361 vs _Ec_Met186) and loss of purine-specific contacts (**Figures 2C** and **S8**). We propound that in WT _Mtb_Rho, NTP hydrolysis is slowed by the Met495 side-chain because its greater-than-leucine bulk hinders changes in the allosteric network that connects the Walker motifs to the RNA-binding Q- and R-loops (**Figures S9** and **S10**). Poorly bound pyrimidine triphosphates may thus dissociate before hydrolysis whereas purine triphosphates are sufficiently stabilized by the network of _Mtb_Rho interactions to the base moiety (**Figures 2C**). We speculate that steric hindrance is diminished by the M495L mutation to the extent that hydrolysis of pyrimidine triphosphates becomes faster than their dissociation from the ATPase pockets.

Taken together, these data illustrate how the resistance of _Mtb_Rho to BCM has been acquired at the expense of enzymatic proficiency. A similar evolutionary tradeoff has been observed previously for BCM-resistant *E. coli* strains bearing rare Rho mutations^32^. These observations and the very high level of phyletic conservation of the BCM binding cavity^4^ support that the configuration of the cavity is substantially constrained by Rho function, thereby making it an attractive target for future drug development.

## CONCLUSION

Upon solving the first structure of a Rho factor from a Gram-positive bacterium (**Figure 1**), we illustrated the general principles and lineage-specific variations that govern RDTT across the bacterial kingdom. Furthermore, we elucidated the molecular mechanism of resistance to bicyclomycin displayed by *M. tuberculosis* and, most likely, by Corynobacteriales relatives (**Figure S11**). This information may be used for future drug development using rational, structure-based approaches. Despite these notable achievements, further work will be needed to fully unravel the role(s) and mechanism of action of the poorly conserved, yet functionally important NID of _Mtb_Rho. Identification of cognate RNA ligands will likely constitute a critical step towards this goal.

## EXPERIMENTAL PROCEDURES

### Materials

Chemicals and enzymes were purchased from Sigma-Aldrich and New England Biolabs, respectively. Bicyclomycin (BCM) was purchased from Santa Cruz Biotechnology. Radionucleotides were from PerkinElmer. DNA templates for *in vitro* transcriptions were prepared by standard PCR procedures, as described previously^33^. RNA substrates were obtained by *in vitro* transcription of PCR amplicons with T7 RNA polymerase and purified by denaturing polyacrylamide gel electrophoresis (PAGE), as described^34^. Plasmid for overexpression of the M495L mutant was prepared by Quickchange (Stratagene) mutagenesis of the pET28b-MtbRho plasmid encoding WT _Mtb_Rho (kindly provided by Dr. Rajan Sen, Hyderabad, India).

### Expression and purification of MtbRho

WT _Mtb_Rho and the M495L mutant were overexpressed in Rosetta 2(DE3) cells (Merck-Millipore) harboring the appropriate pET28b derivative and purified following published protocols^35^ with minor modifications. Briefly, crude protein lysates were fractionated by polymin-P and ammonium sulfate precipitations before purification by affinity chromatography on a HisTrap FF column, cation exchange chromatography on a HiTrap SP sepharose, and gel filtration chromatography on a HiLoad 16/600 Superdex 200 column (all columns from GE Healthcare). Purified _Mtb_Rho proteins were used directly for Cryo-EM or stored at −20°C as micromolar hexamer solutions in storage buffer (100 mM KCl, 10 mM Tris-HCl, pH 7.9, 0.1 mM EDTA, 0.1 mM DTT, 50% glycerol) for biochemical assays.

### Cryo-EM sample preparation and data collection

The _Mtb_Rho-ATP-DNA complex sample was prepared by mixing 0.6 mg/mL of freshly purified WT _Mtb_Rho with 10 μM dC_20_ oligonucleotide and 1mM ATP^26^ in cryoEM buffer (150 mM NaCl, 5 mM MgCl_2_, 10 mM Tris-HCl, pH 7.6). Three microliters of the mixture were applied to glow-discharged Lacey 300 mesh copper grids (Ted Pella Inc.), blotted for 3-4 s, and then flash frozen in liquid ethane using the semi-automated plunge freezing device Vitrobot Mark IV (ThermoFisher Scientific) maintained at 100% relative humidity and 22°C. Preliminary images were recorded using a JEOL 2200 FS electron microscope operating at 200 kV in zero-energy-loss mode with a slit width of 20 eV and equipped with a 4k × 4k slow-scan CCD camera (Gatan inc.). High resolution data were collected at the EMBL cryo-EM core facility (Heidelberg, Germany) with a Titan Krios S-FEG transmission electron microscope (ThermoFisher Scientific) operating at 300 kV at a nominal magnification of X 165,000 under low-dose conditions with defocus values ranging from −0.8 to −2 μm, using the SerialEM automated acquisition software^36^. Movies were recorded with a K2-Summit direct electron detector (Gatan Inc.) configured in counting mode, mounted on a Gatan Quantum 967 LS energy filter using a 20 eV slit width in zero-loss mode. The scheme of image recording was made with 7 images per position plus 1 middle, and as a series of 40 frames per movie, with 1.2 e^−^/Å^2^ per frame, therefore giving a total accumulated dose of 48 e^−^/Å^2^, with a corresponding pixel size of 0.81 Å/pix.

### Image Processing and high-resolution 3D reconstruction

A total of 10,888 movies were recorded. The frames of each movie were computationally dose-weighted and corrected for drift and beam-induced movement in RELION-3.1.0^37^ own implementation. The contrast transfer function (CTF) of each micrograph was determined using Gctf-v1.18 program^38^. A total of 2,217,252 particles were automatically picked with RELION’s autopicking procedure. Briefly, a template-free autopicking procedure based on a Laplacian-of-Gaussian (LoG) filter allowed to select and extract a set of particles from an initial small set of images and to compute corresponding 2D class averages. The best 2D class averages were then used as references in a second round of autopicking in order to optimize parameters. Once determined, the autopicking procedure was applied to all images. Particles were initially extracted with a box size of 320 X 320 pixels and binned to obtain a pixel size of 3.25 Å /pix, and submitted to 2D classification. The best 2D class averages allowed to select 1,390,182 particles. A 3D classification with 4 classes was then carried out. Two classes corresponding to 986,385 particles were selected and particles re-extracted with a box size of 400 X 400 pixels and binned at 1.49 Å /Pix. After two rounds of 3D auto-refinement without symmetry, a first density map at 3.97 Å was obtained. We then proceeded with per-particle defocus estimation, beam-tilt estimation plus Bayesian polishing. The final 3D refinement allowed to compute a map at 3.32 Å resolution (FSC = 0.143) after post-processing. Figures were prepared using Chimera^39^.

### Structure refinement

The map coefficients were converted into structure factors using the corresponding Phenix cryo-EM tool^40^, and molecular replacement was performed with Molrep^41^, using one of the monomers of _Ec_Rho (chain C of PDB entry 1PV4) as search model. Four of the six chains were immediately placed and the remaining two (the outer ones of the open ring) were added in subsequent runs. The electron density corresponding to these outer chains (A and F) was overall weaker and less clear than that of the inner chains. The homohexameric structure was initially refined by Refmac^42^ using the maximum-likelihood method, and subsequently with the Phenix real-space refinement tool^43^. Non-crystallographic symmetry, secondary structure, Ramachandran and rotamer restraints were used. The refinement cycles were alternated with extensive manual model-building, fitting and validation in Coot^44^. Six ATP molecules were fitted into clearly visible electron density, but no density imputable to nucleic acid could be identified. Additional density found near the β and γ-phosphates in five out of the six chains was assigned to catalytic Mg^2+^ ions coordinated to β and γ-phosphate oxygens. No electron density corresponding to residues before Val222 and after Ser592 (before Val223 and after Val589 for chain F) was visible. Gaps in the density meant that residues after Lys277 and before Phe288 in chain C could not be modeled (corresponding residues for the other chains are A: 278-287, B: 276-285, D: 278-286, E: 276-286, F: 279-287). The final refinement run gave a model-to-map fit (CC_mask) of 0.7753, Molprobity all-atom clashscore of 11.32, 0.05% outliers in the Ramachandran plot (93.47% residues in favored and 6.49% in allowed geometries) and 0.3% sidechain outliers. The cryo-EM data and the model were deposited as entries EMD-12701 and PDB 7OQH.

### NTPase assays

NTP hydrolysis activities were determined with a thin layer chromatography (TLC) assay, as described previously^45^. For each NTP, a mixture of the ‘cold’ NTP and matching α[^32^P]-NTP (or γ[^32^P]-NTP) was used. Reaction mixtures contained 20 nM Rho hexamers, 1 mM of the NTP/[^32^P]-NTP mixture, and 10 μM poly[rC] in NTPase buffer (50 mM KCl, 1 mM MgCl_2_, 20 mM HEPES, pH 7.5, 0.1 mM EDTA, and 0.1 mM DTT) and were incubated at 37°C. Reaction aliquots were withdrawn at various times, quenched with four volumes of 0.5 M EDTA, and stored on ice. Once all collected, the aliquots were spotted on a PEI-cellulose TLC plate (Sigma-Aldrich) and developed with 0.35 M potassium phosphate buffer (pH 7.5). Plates were dried and analysed by phosphorimaging with a Typhoon FLA 9500 imager (GE Healthcare).

### Duplex unwinding assays

RNA:DNA duplexes were prepared by mixing 5 pmoles of ^32^P-end labelled and 30 pmoles of unlabelled RNA strand with 60 pmoles of complementary DNA oligonucleotide (see **Table S2** for sequences) in helicase buffer (150 mM potassium acetate, 20 mM HEPES, pH 7.5, 0.1 mM EDTA). Mixtures were heated at 90°C and slowly cooled to 20°C. Duplexes were then purified by native 6% PAGE and stored in helicase buffer at −20°C before use. Helicase reactions were performed as described previously^46^ with minor modifications. Briefly, 5 nM ^32^P-labelled RNA:DNA duplex were mixed with 20 nM Rho hexamers in helicase buffer supplemented with 0.1 mg/mL BSA and incubated for 3 min at 30°C. The helicase reaction was initiated by addition of a mix containing MgCl_2_ and ATP (1 mM, final concentrations) and an excess of Trap oligonucleotide (400 nM final concentration) before further incubation at 30°C. Reaction aliquots were taken at various times, mixed with two volumes of quench buffer (30 mM EDTA, 0.75% SDS, 150 mM sodium acetate, 6 % Ficoll-400), and loaded on 8% PAGE gels containing 0.5% SDS. Gels were analysed by Typhoon phosphorimaging, as described^46^.

### Transcription termination experiments

Standard transcription termination experiments were performed as described previously^6,27,33^. Briefly, transcription reactions were assembled by mixing DNA template (0.1 pmol), *E. coli* RNAP (0.3 pmol), Rho (0 or 1.4 pmol), Superase-In (0.5U/μL; Ambion) and BCM (0 to 15 nmol, Santa) in 18 μL of transcription buffer (40 mM Tris-HCl, pH 8, 5 mM MgCl_2_, 1.5 mM DTT, and 100 mM KCl). Mixtures were incubated for 10 min at 37°C before addition of 2 μL of initiation solution (250 μg/mL rifampicin, 2 mM ATP, GTP, and CTP, 0.2 mM UTP, and 2.5 μCi/μL α[^32^P]UTP in transcription buffer). After 20 min of incubation at 37°C, reactions were stopped upon addition of 4 μL of EDTA (0.5 M), 6 μL of tRNA (0.25mg/mL), and 80 μL of sodium acetate (0.42 M) followed by ethanol precipitation. Reaction pellets were resuspended in loading buffer (95% formamide; 5mM EDTA) and analyzed by denaturing 7% PAGE and Typhoon phosphorimaging. Transcript sizes were estimated by comparing gel band migrations with those of control RNA/DNA species and used for rough α[^32^P]U content normalization of band intensities (Rabhi et al., 2011). The normalized band intensities were used to estimate termination efficiencies from which IC_50_ values for BCM inhibition were deduced, as described previously^6,27^.

## Acknowledgements

We warmly thank Annie Schwartz for her help with biochemical experiments. This work benefited from access to the cryo-EM facility of the European Molecular Biology Laboratory (EMBL) in Heidelberg with support from the iNEXT-Discovery program (project #871037) funded by the Horizon 2020 framework of the European Commission. The work was supported by grants from the French Agence Nationale de la Recherche (ANR-15-CE11-0024-01 to EM and ANR-15-CE11-0024-02 to MB), a sabbatical research fellowship from LE STUDIUM Loire Valley Institute for Advanced Studies (Marie Sklodowska-Curie 665790) to E.S., and a doctoral fellowship from Région Centre-Val de Loire to I.S. CBS is a member of the French Infrastructure for Integrated Structural Biology (FRISBI) supported by Agence Nationale de la Recherche (ANR-10-INBS-05).

