## Supplementary Information for "Cryo-EM structure of the transcription termination factor Rho from *Mycobacterium tuberculosis* reveals mechanism of resistance to bicyclomycin"

### Crystallization trials with $_{Mtb}$Rho

We performed extensive crystallization trials with the purified WT $_{Mtb}$Rho and T501K mutant. The attempts involved the protein unliganded, as well as complexed with various RNA and DNA oligonucleotides, and ATP analogues[1,2]. None of these trials produced crystals.

We next optimised the oligonucleotide/ cofactor combination using the thermal shift assay method[3,4]. The thermostability ($T_m$) of the protein was measured in the apo- form and upon mixing with various potential ligands. ADP.$BeF_3$ and ADP.$AlF_4$, combined with oligonucleotides $dC_{20}$, $dC_{34}$, and 5'r(UUCCCCCCCC), all led to an increase in thermostability[5] and crystallization screening experiments with most of these combinations were conducted, but still without success.

We thus decided to perform *in situ* random proteolysis, with the addition of trypsin or chymotrypsin directly into crystallisation drops[6,7]. Crystals were obtained with the T501K mutant incubated with 5'r(UUCCCCCCCC)/ ADP.$BeF_3$ and trypsin (1.5 mg/mL original stock, added at a 1:10000(v/v) dilution to T501K), in 13% PEG 3350, 0.2 M $NaSO_4$ and 0.1 M Bis-Tris propane pH 7.5. These crystals, however, only diffracted to ca. 6 Å at best.

Since a crystal structure could not be determined with such crystals, several of them were pooled together, washed and dissolved for analysis by high-resolution mass spectroscopy (at the MS core facility of CBM). Several T501K proteolytic fragments were identified reproducibly in the dissolved material, none of them likely to contain the NID. We surmised that these fragments may retain both their folding and their mutual interactions. This would explain that they could form crystals at all and, more significantly, that the molecular replacement solution given by the low resolution crystallographic data indicated an expected hexameric conformation. The likely absence of the NID of $_{Mtb}$Rho from the dissolved crystalline material would confirm its deleterious effect for crystallization.

### Characterization of the open Rho hexamers

In order to compare the 3D ring structures of $_{Ec}$Rho and $_{Mtb}$Rho, two distance parameters, the 'ring opening' and 'rise' between protomers at the gap (**Figure S5**), were defined according to the following protocol. For each open ring structure of $_{Mtb}$Rho (this work) and $_{Ec}$Rho in isolation (PDB 1PVO, 1PV4, 1XPO, 6WA8) or in complex with RNAP (PDB 6XAS, 6Z9P, 7ADB), the MatchMaker module of Chimera was used to superimpose the C-terminal domain of the protomer 1 (residues 1-129 for $_{Ec}$Rho and 1-306 for $_{Mtb}$Rho were omitted) on protomer 1 of the reference, i.e. closed $_{Ec}$Rho ring structure (PDB 3ICE) (see **Figure S7** for protomer numbering). Then, the center of mass (COM) of the C-terminal domain of each Rho protomer was calculated using the CALCOM server (http://bioinformatica.isa.cnr.it/CALCOM) and displayed as a 2 Å radius sphere (**Figure S7**). The 'ring opening' parameter represents the distance in Angstroms between the COMs of protomers 1 and 6 at the gap (**Figure S7,** dash line) minus the distance between COMs 1 and 6 in the 3ICE reference. The 'rise' parameter is defined as the shortest distance between the COM of protomer 6 of the structure of interest and a plane passing through the six COMs of the 3ICE reference.

**Table S1:** Cryo-EM data collection, refinement and validation statistics

| | Rho-ATP-ADN (EMDB-12701) (PDB 7OQH) |
|---|---|
| ***Data collection and processing*** | |
| Magnification | 165,000 |
| Voltage (kV) | 300 |
| Electron exposure (e–/Å$^2$) | 48.07 |
| Defocus range (μm) | -0.8 to -2 |
| Pixel size (Å) | 0.81 |
| Symmetry imposed | C1 |
| Initial particle images (no.) | 2,217,252 |
| Final particle images (no.) | 986,385 |
| Map resolution (Å) | 3.32 |
| FSC threshold | 0.143 |
| Map resolution range (Å) | 3.1-4.7 |
| Map sharpening B factor (Å$^2$) | -107.639 |
| ***Refinement*** | |
| Model resolution (Å) | 3.3 |
| FSC threshold | 0.143 |
| Model composition | |
| Non-hydrogen atoms | 16,824 |
| Protein residues | 2,169 |
| Ligand atoms (ATP) | 186 (6 ATP) |
| Ligand atoms ($Mg^{2+}$) | 5 |
| B factors (Å$^2$) | |
| Protein | 36.52 |
| Ligand (ATP) | 38.28 |
| Ligand ($Mg^{2+}$) | 28.89 |
| R.m.s. deviations | |
| Bond lengths (Å) | 0.004 |
| Bond angles (°) | 0.616 |
| ***Validation*** | |
| MolProbity score | 2.00 |
| Clashscore | 11.3 |
| Poor rotamers (%) | 0.3 |
| Ramachandran plot | |
| Favored (%) | 93.47 |
| Allowed (%) | 6.49 |
| Disallowed (%) | 0.05 |

| **Table S2**: Nucleic acid sequences used in the duplex unwinding assay | |
|---|---|
| **RNA strand** | 5’r(GGACUUCUCCUCUGUCUCCUUCUUCCUUCUCCUCUGUCUCCUUCUUCCUCGACCUAUUGAGUUUGAAUUUAUCGAUGGUAUCAGAUCUGGAUCCUCGAGAAGCUGCGGGUACCGAGCUCGAAUUCAUCG) |
| **DNA strand** | 5’d(CGATGAATTCGAGCTCGGTACCCGCAGCTTCTCGAGGATCCAGATCTGATACCATCG) |
| **Trap oligo** | 5’d(CGATGGTATCAGATCTGGATCCTCGAGAAGCTGCGGGTACCGAGCTCGAATTCATCG) |

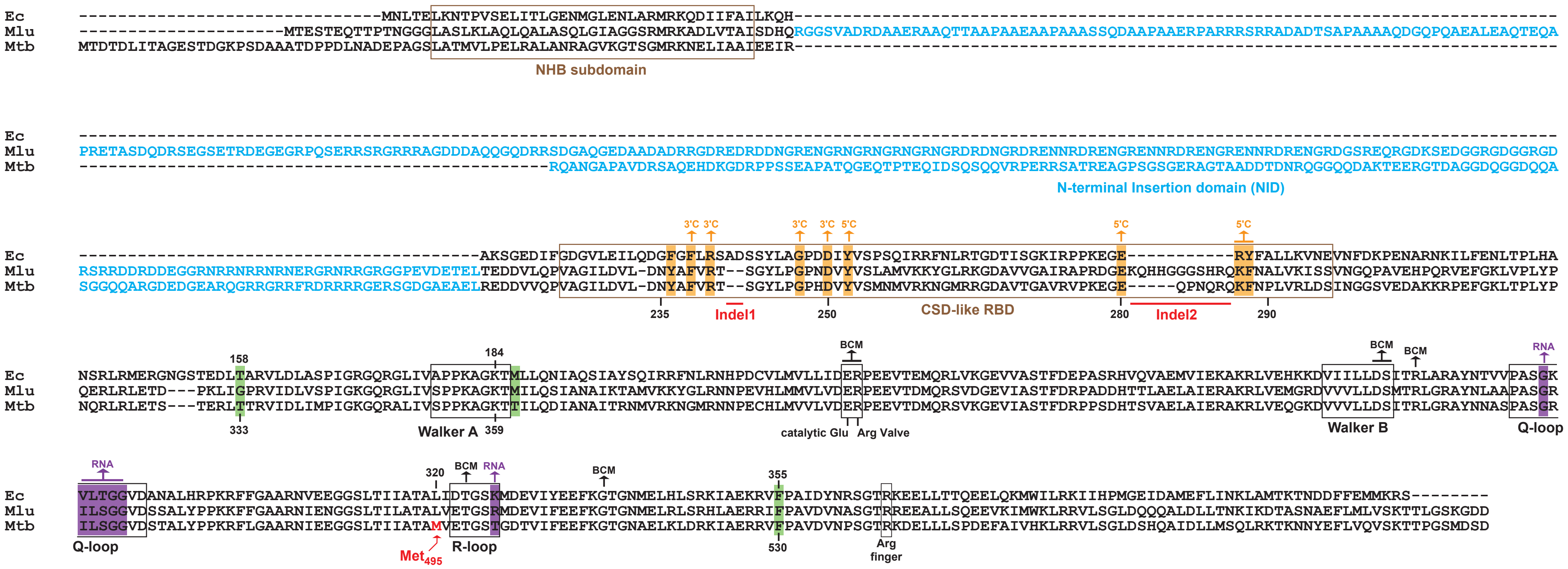


**Supplementary Figure 1:** Alignment of the Rho sequences from *E. coli* (Ec), *M. luteus* (Mlu), and *M. tuberculosis* (Mtb). Positions corresponding to PBS and SBS residues in $_{Ec}$Rho are boxed in orange and purple, respectively. Residues contacting the base moiety of ATP are boxed in green.

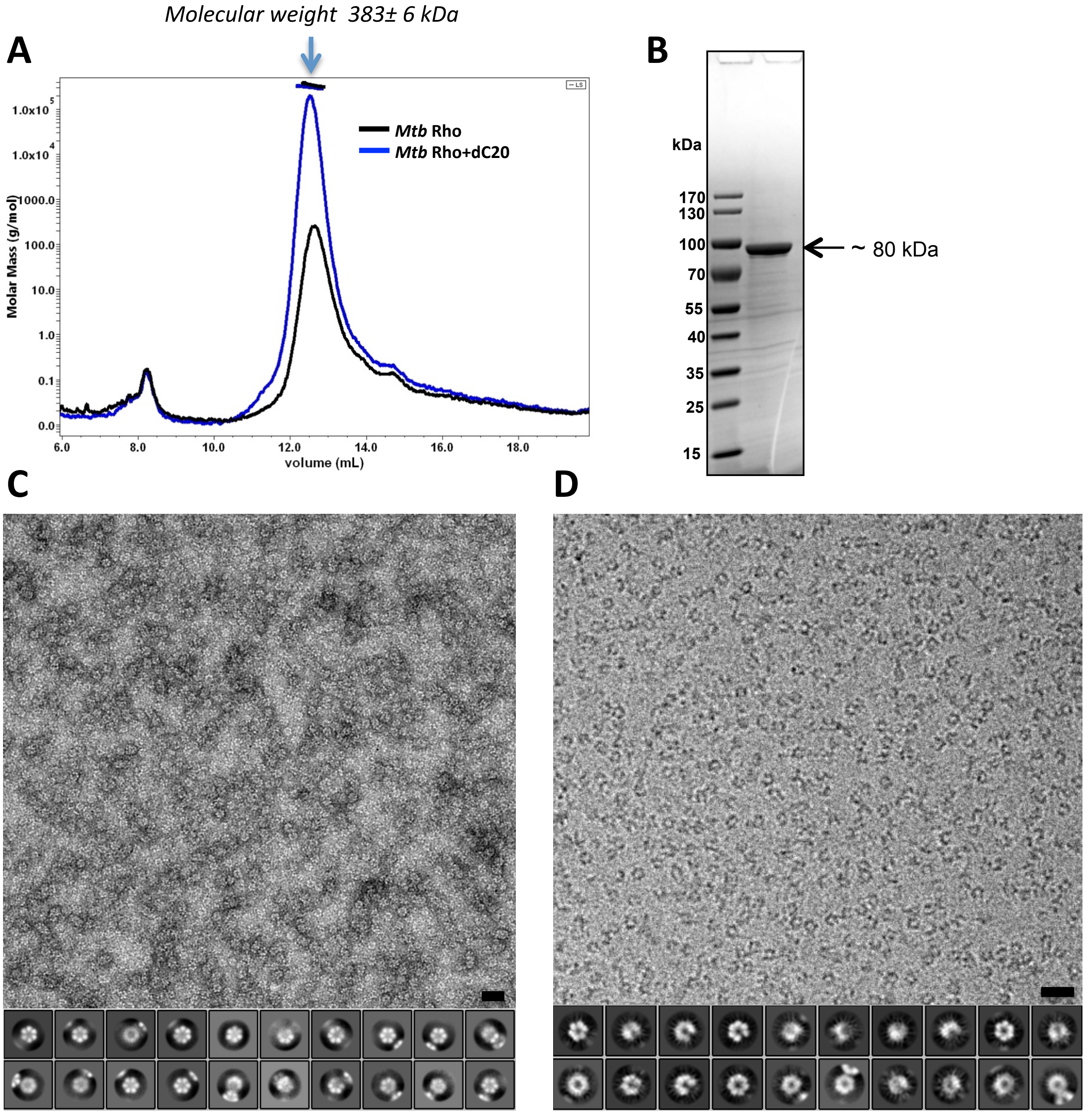


**Supplementary Figure 2 :** Purification and preliminary characterization of $_{Mtb}$Rho**. (A)** Size Exclusion Chomatography - Multi-Angle Light Scattering (SEC-MALS) profile. The highest peak located between 12.0 to 14.0 mL corresponds to a molecular weight of 383± 6 kDa. The theoretical molecular weight of $_{Mtb}$Rho monomer is about 65 kDa. So this peak corresponds to a hexameric organisation of $_{Mtb}$Rho. **(B)** Sodium dodecyl sulphate–polyacrylamide gel electrophoresis (SDS-PAGE) of the highest peak fraction. It reveals a unique band at about 80 kDa. The mass shift observed between the apparent weight and the theoretical molecular weight of $_{Mtb}$Rho has already been observed and due to an abnormally slow migration. **(C)** Negative stain image of $_{Mtb}$Rho complex in presence of Mg-ATP and dC20 ligands from in-house JEOL2200FS microscope and corresponding 2D class averages. $_{Mtb}$Rho particles are homogeneous, showing and confirming the hexameric organization of $_{Mtb}$Rho, where only top views are visible when $_{Mtb}$Rho partices are bound to the carbon film. **(D)** Cryo-EM image of $_{Mtb}$Rho complex in presence of Mg-ATP and dC20 ligands from in-house JEOL2200FS microscope and corresponding 2D class averages. Image and 2D class-averages of frozen-hydrated $_{Mtb}$Rho complexes reveal various orientations of particles in ice. So $_{Mtb}$Rho is suitable for high resolution acquisition. Scale bars, 60 nm.

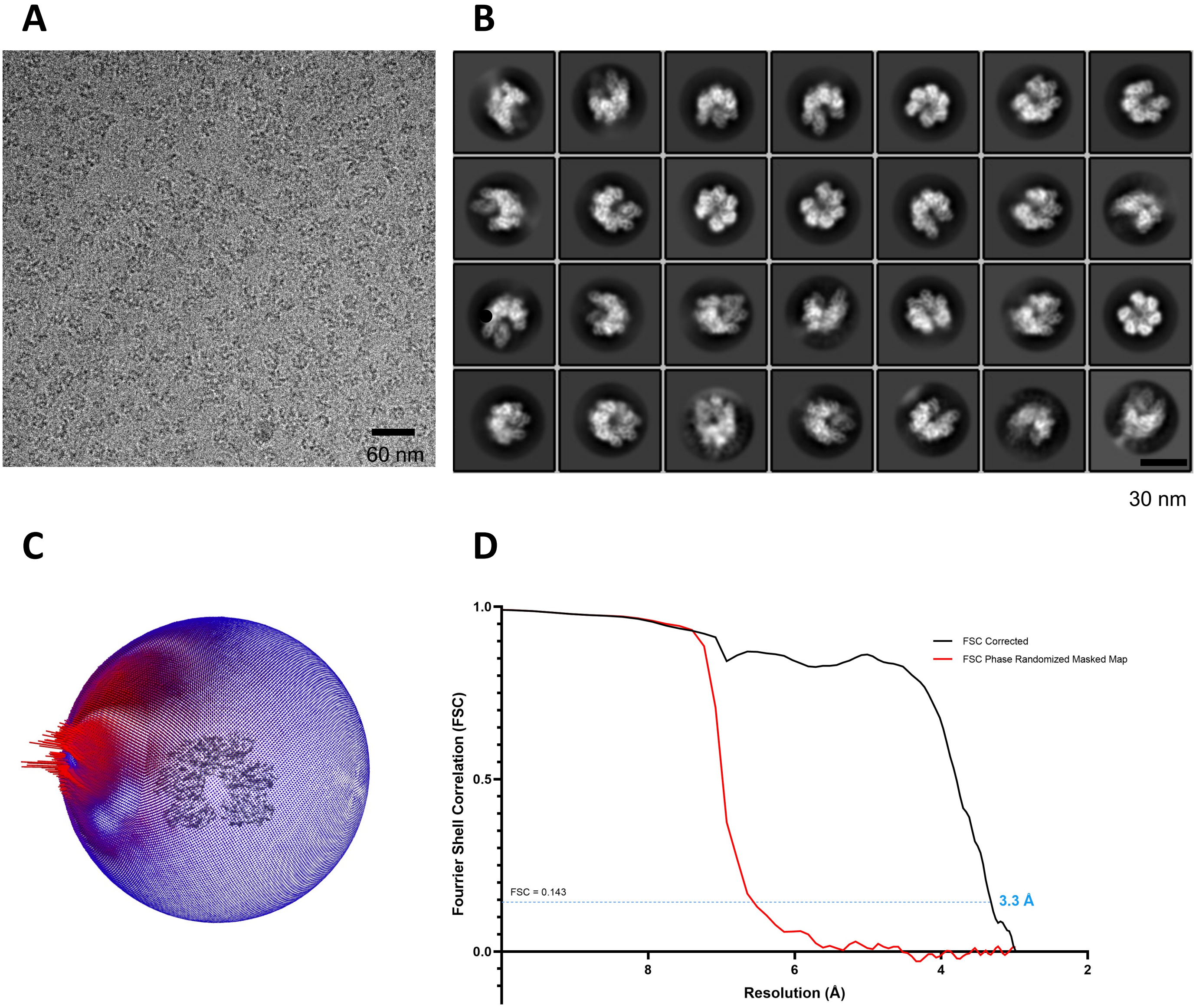


**Supplementary Figure 3 :** Cryo-EM analysis of $_{Mtb}$Rho transcription factor complex. **(A)** Representative motion-corrected electron micrograph embedded in vitreous ice (0.81 Å per pixel, 1.86 um defocus) of the $_{Mtb}$Rho dataset (scale bar, 60 nm). This image has been lowpass-filtered to 1 nm for better visibility. **(B)** Representative reference-free two-dimensional class averages of $_{Mtb}$Rho complex (scale bar, 30 nm), from Relion two-dimensional classification. These 2D class averages show secondary structure elements. **(C)** Euler angle distribution plots of $_{Mtb}$Rho particles, reflecting the initial angular distribution. The number of particles with respective orientations are represented by length and colored cylinders, ranging from blue to red. **(D)** High resolution refinement : the Fourier Shell Correlation (FSC) curves between independently refined half-maps at the final stage of the processing indicates an average resolution of 3.3 Å according to the FSC=0.143 criterion.

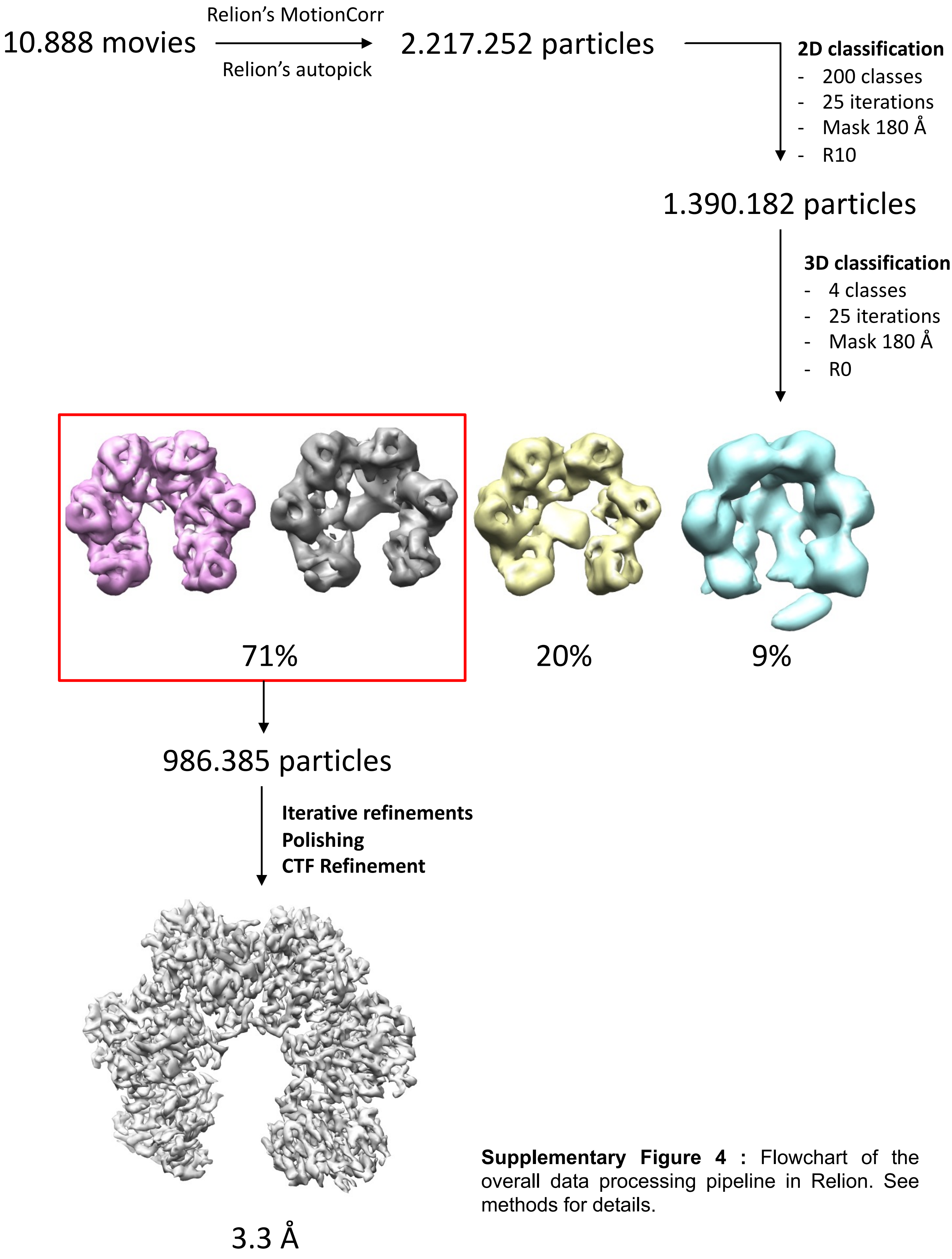


**Supplementary Figure 4 :** Flowchart of the overall data processing pipeline in Relion. See methods for details.

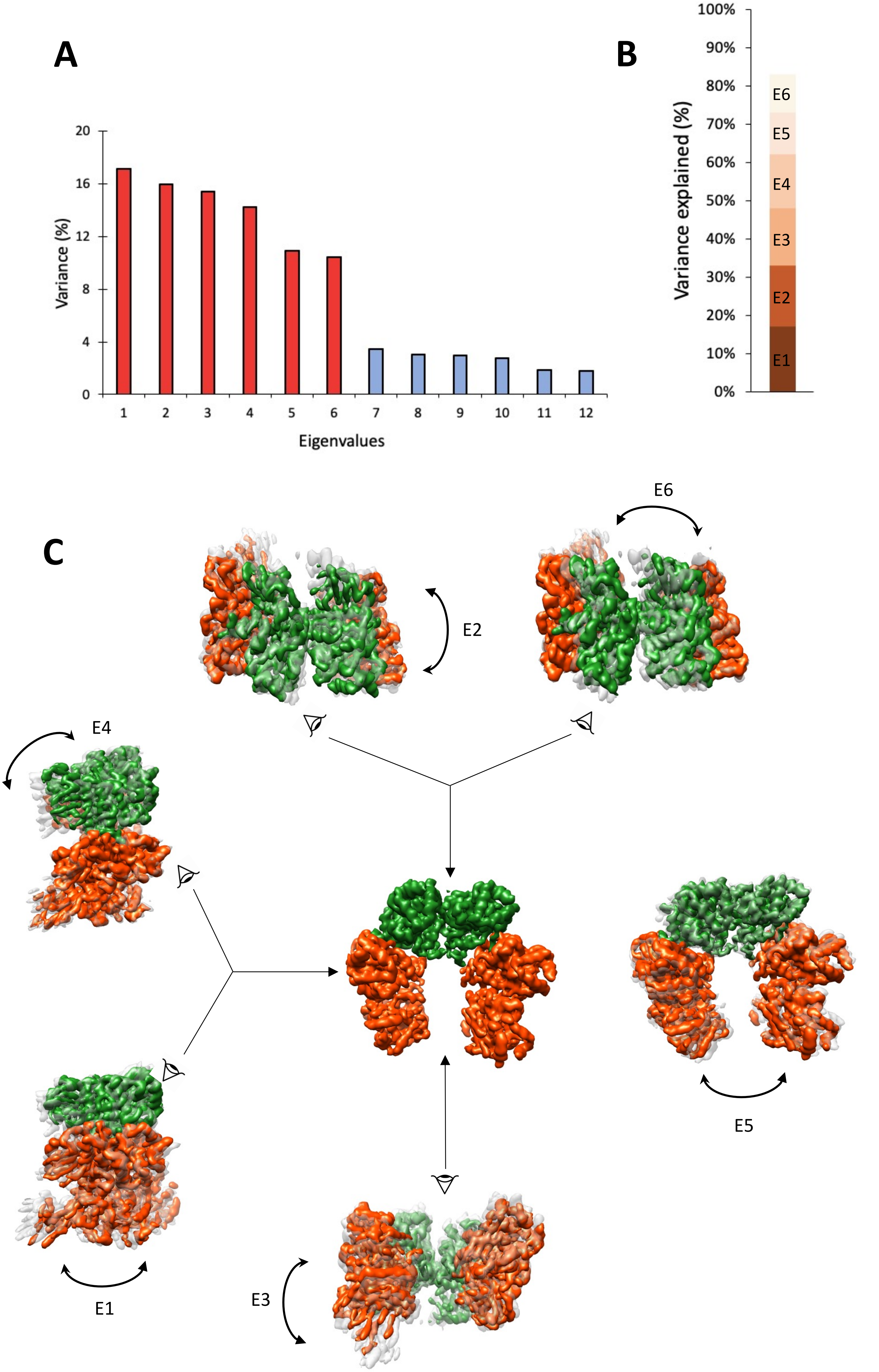


**Supplementary Figure 5 :** Structural flexibility and dynamics of the $_{Mtb}$Rho hexamer. We observe interdomain and intersubunit movements in the structure. **(A)** Contribution of each of 12 eigenvectors to the variance of the final cryo-EM map. **(B)** Eigenvectors 1 to 6 correspond to most (83%) of the variance. **(C)** The $_{Mtb}$Rho consensus map (3.4 Å) used for multibody 3D-refinement is shown in the middle, with two bodies colored interdomain dynamics (in gray). The most resolved domains are in green (chains C/D) while others are in orange/red. Maps corresponding to the first 6 eigenvectors are shown. E4 is part of E1 while E6 is part of E2.

.

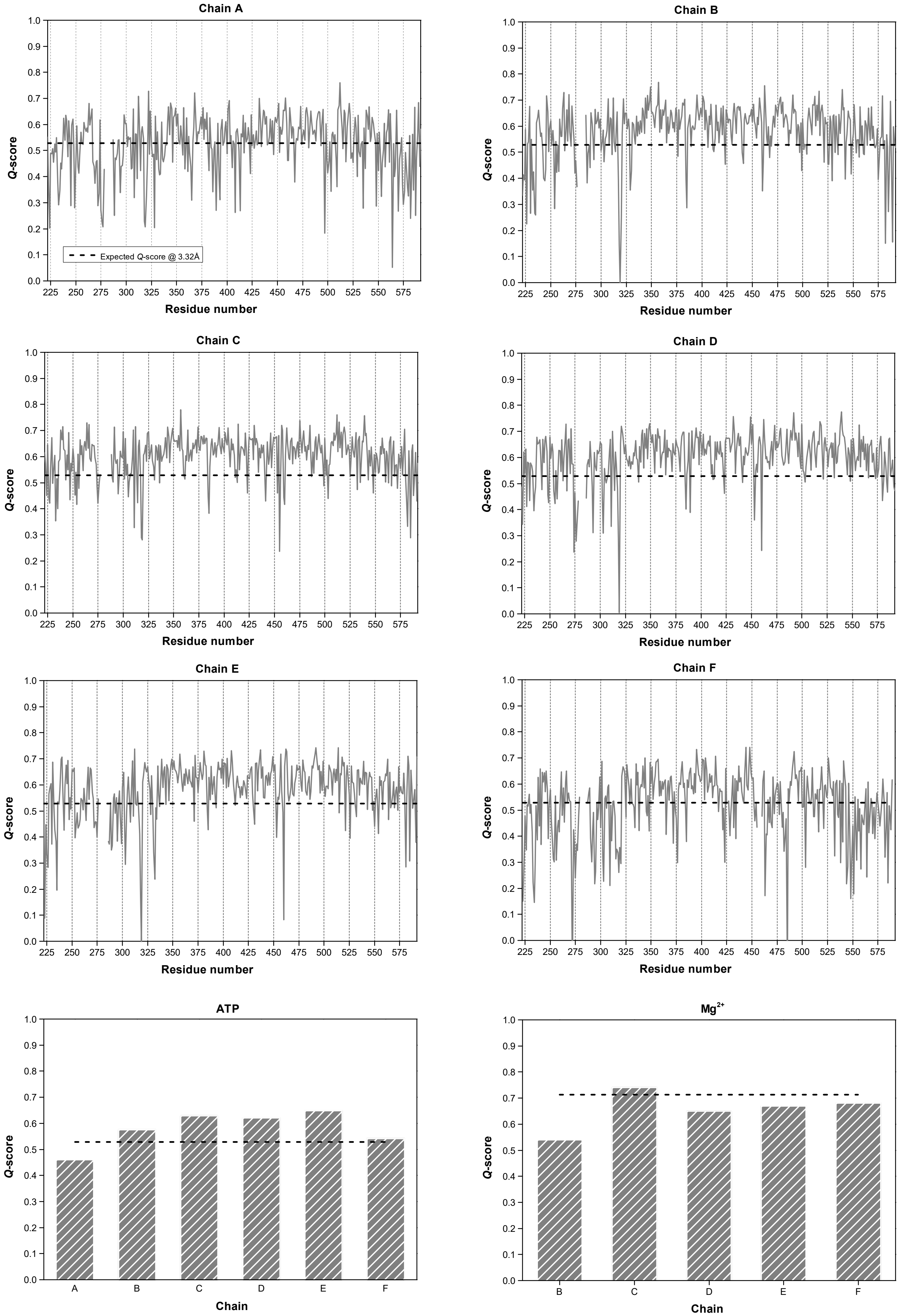


**Supplementary Figure 6:** Residue resolvability in cryo-EM map. Q scores were calculated with the MapQ plugin of UCSP Chimera. Dashed lines represent the average Q-score expected at 3.32 Å resolution.

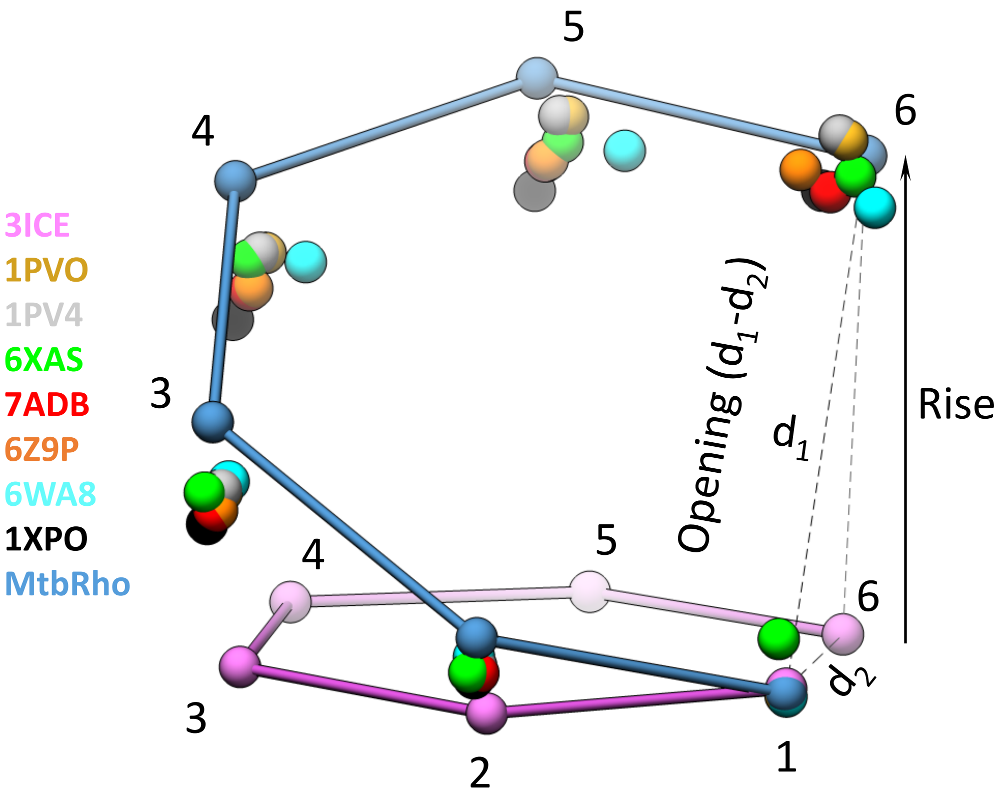


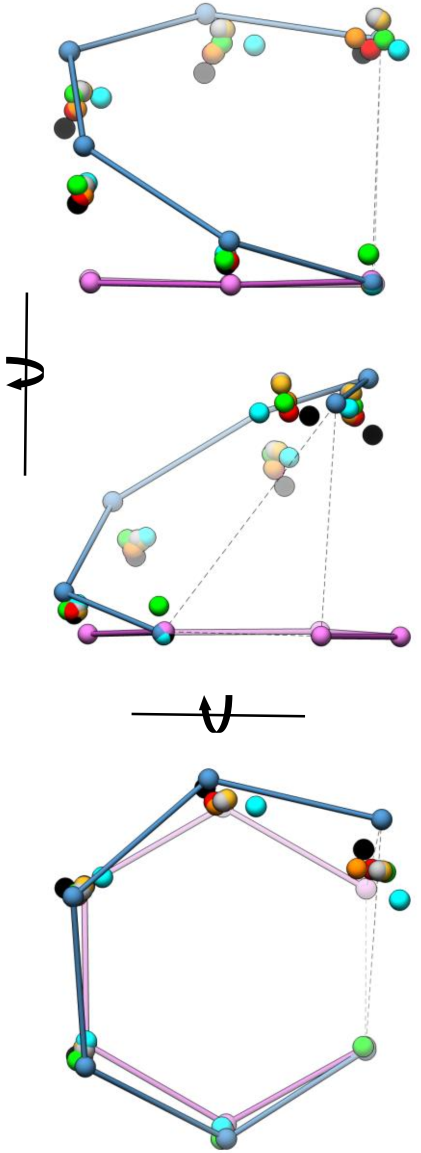

| | PDB# | Opening (Å) | Rise (Å) |
|---|---|---|---|
| $_{Ec}$Rho (X-ray) | 3ICE | 0 | 0 |
| | 1PVO | 25 | 50 |
| | 1PV4 | 24 | 51 |
| | 1XPO | 22 | 45 |
| $_{Ec}$[Rho:RNAP] complexes (cryoEM) | 6XAS | 17 | 43 |
| | 7ADB | 20 | 46 |
| | 6Z9P | 22 | 49 |
| | 6WA8 | 18 | 45 |
| $_{Mtb}$Rho (this work) | 7OQH | 27 | 47 |

**Supplementary Figure 7:** Rho ring parameters.

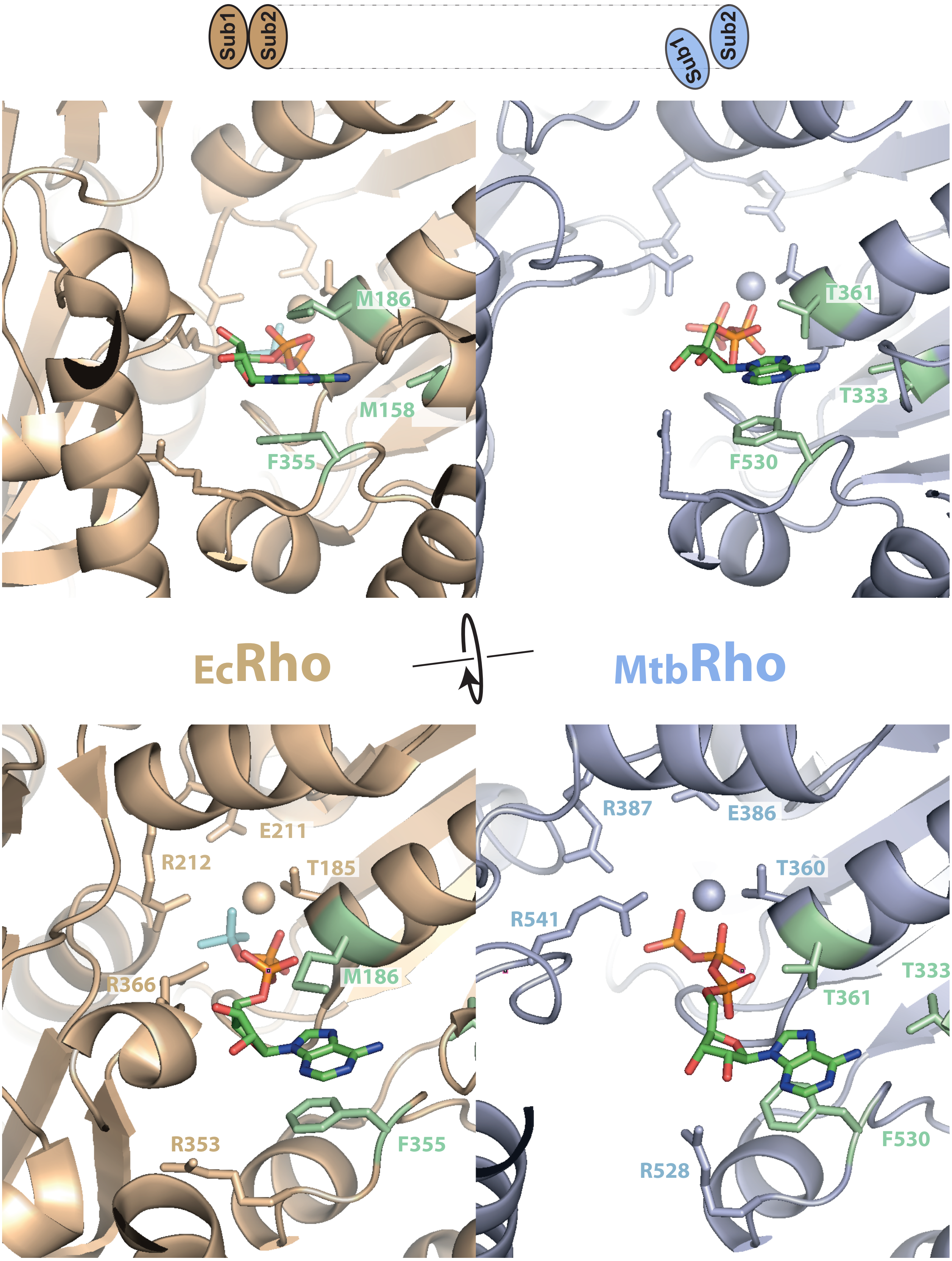


**Supplementary Figure 8:** Organization of the ATPase pockets in the closed $_{Ec}$Rho hexamer (PDB 5JJI) and open $_{Mtb}$Rho hexamer (this work). The B/C subunit interface of $_{Ec}$Rho (interface in a productive ATPase conformation in the assymetric closed hexamer) and C/D subunit interface of $_{Mtb}$Rho (best resolved interface) were used for comparison. The $BeF_3$ ion in the $_{Ec}$Rho structure is represented by cyan sticks. $Mg^{2+}$ ions are shown as spheres. The respective dispositions of subunits at the interfaces are schematically depicted above figures.

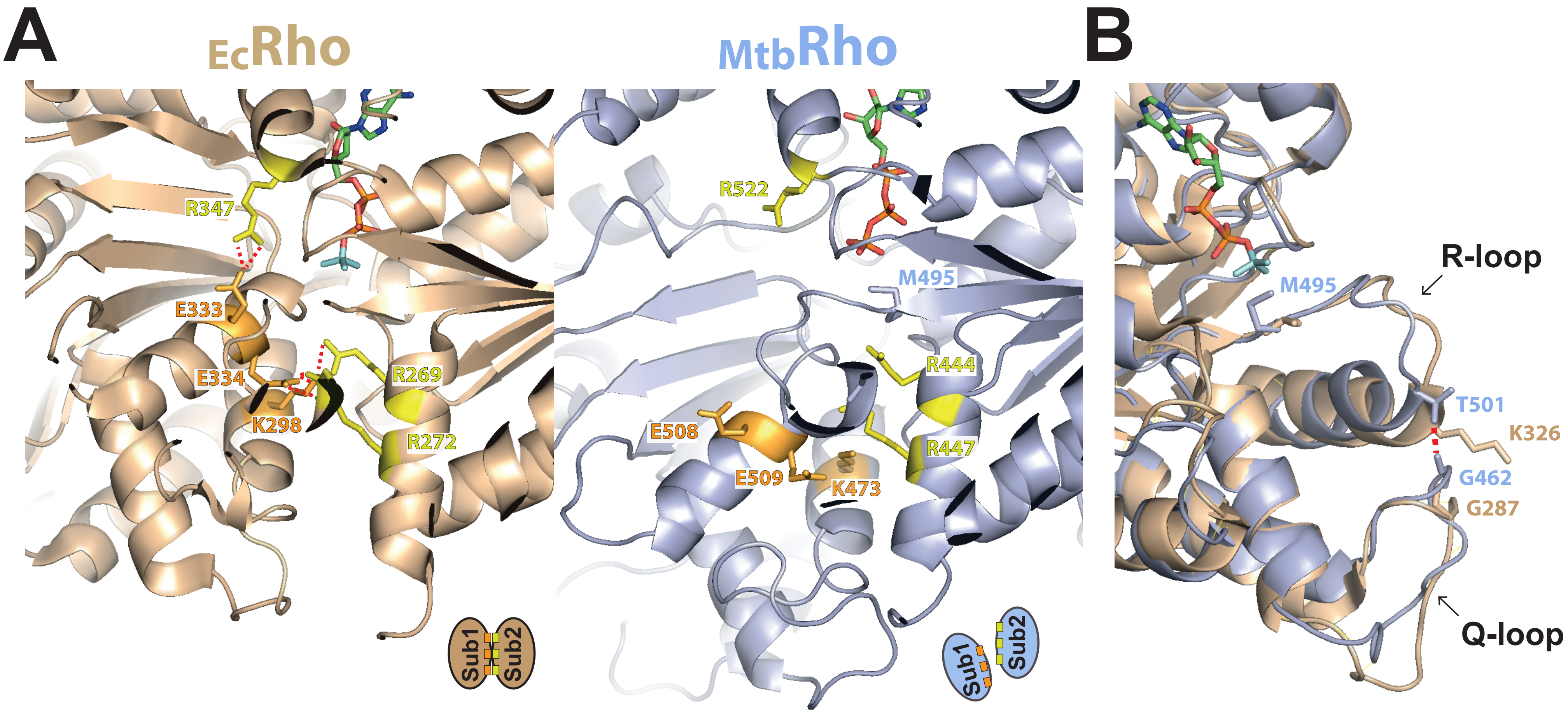


**Supplementary Figure 9:** Organization of the allosteric communication network in the closed $_{Ec}$Rho hexamer (PDB 5JJI) and open $_{Mtb}$Rho hexamer (this work). **(A)** Network contacts between subunits are disrupted in $_{Mtb}$Rho, which is consistent with an unproductive hexamer conformation. **(B)** The Thr501 side-chain of $_{Mtb}$Rho mediates an interaction between the Q- and R-loops. In $_{Ec}$Rho, the corresponding Lys326 side-chain lies in the central channel where it can contact and translocate the RNA chain (Thomsen & Berger, Cell, 2009, 139, 523-24).

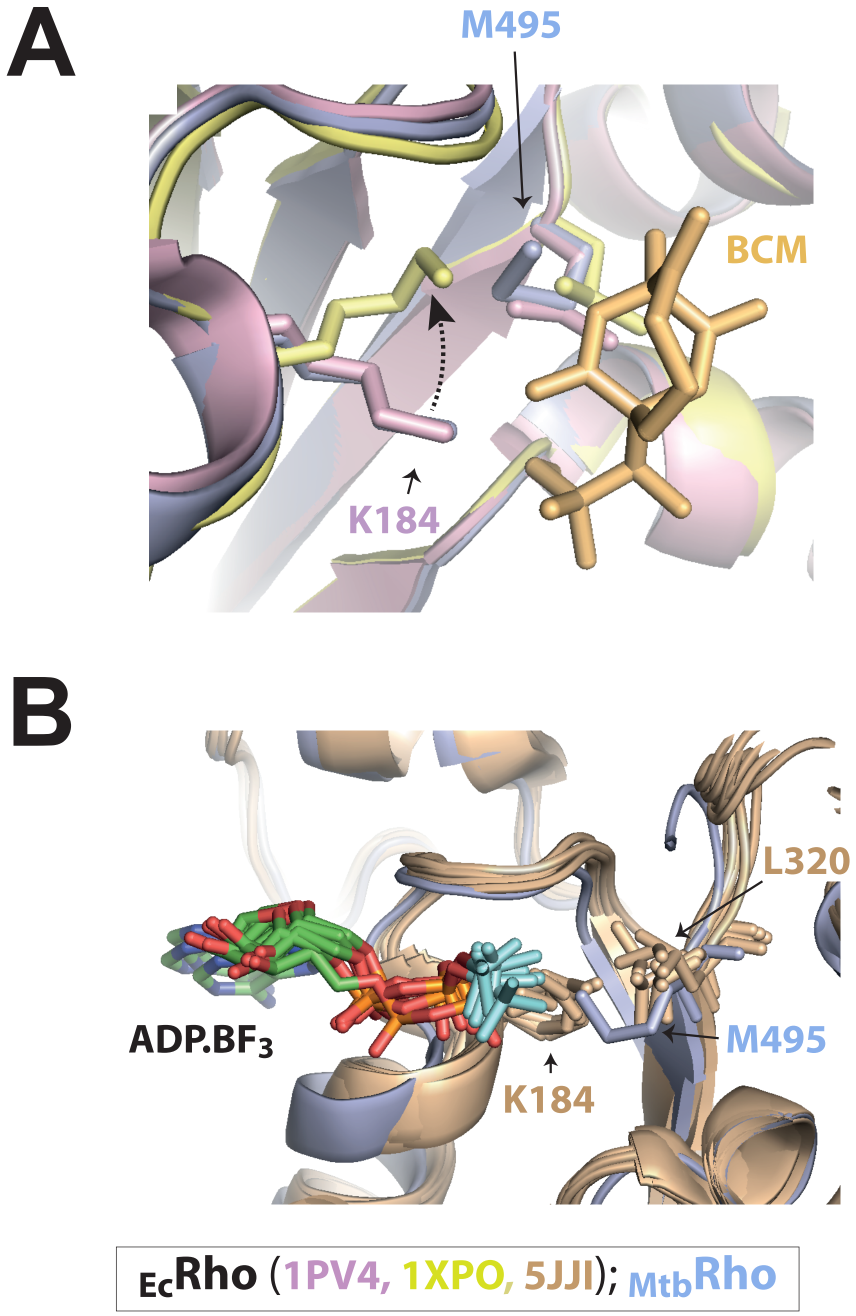


**Supplementary Figure 10:** The bulky $_{Mtb}$Met495 side-chain lies in the path of a mobile lysine from the ATPase Walker A motif ($_{Mtb}$Lys380, i.e. $_{Ec}$Lys184 in $_{Ec}$Rho). **(A)** The $_{Ec}$Lys184 side-chain adopts distinct conformations in BCM-free (in pink) and BCM-bound (in light yellow) $_{Ec}$Rho (protomers C from PDB 1PV4 and 1XP0, respectively). The $_{Mtb}$Rho protomer C is in blue and BCM in light orange. **(B)** Motion of the $_{Ec}$Lys184 side-chain in the assymetric, closed $_{Ec}$Rho hexamer (PDB 5IJJ) as a function of the chemical/conformational state of the ATPase pocket. The six $_{Ec}$Rho protomers (in light brown) have been aligned with $_{Mtb}$Rho protomer C (in blue). The $BeF_3$ ion is shown in cyan sticks.

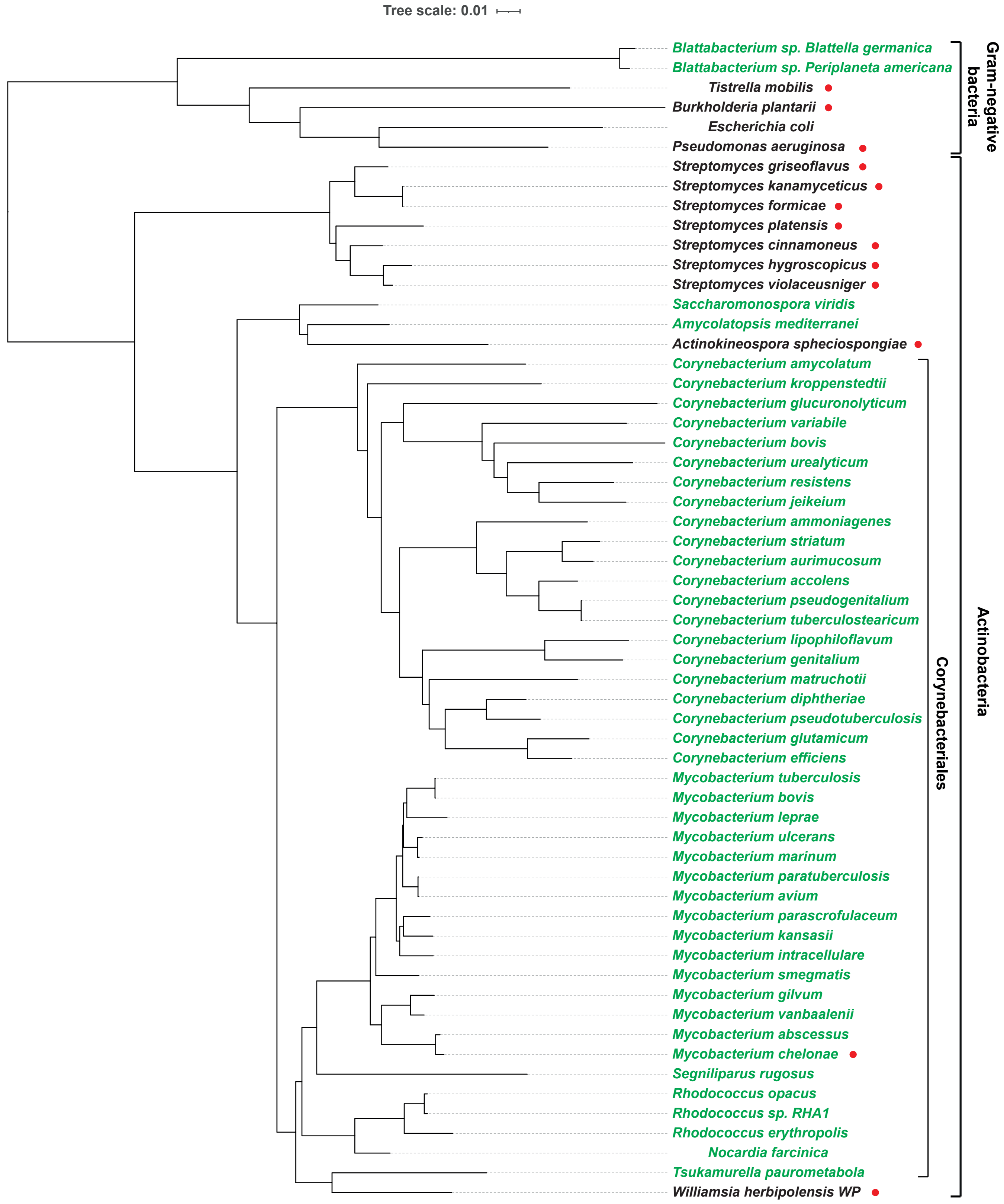


**Supplementary Figure 11:** Phylogenetic distribution of the Leu→Met substitution in the BCM-binding pocket of Rho. Species bearing the Leu→Met mutation are in green. Most belong to the Corynebacteriales order. *Blattabacterium sp.* (Bacteroidetes phylum) are insect endosymbionts. Species bearing the cluster of genes for BCM biosynthesis are identified by a red dot. The unrooted tree has been built with the Seaview 4.6.4 software.
